# Topological phase transitions in functional brain networks

**DOI:** 10.1101/469478

**Authors:** Fernando A. N. Santos, Ernesto P. Raposo, Maurício D. Coutinho-Filho, Mauro Copelli, Cornelis J. Stam, Linda Douw

## Abstract

Functional brain networks are often constructed by quantifying correlations among brain regions. Their topological structure includes nodes, edges, triangles and even higher-dimensional objects. Topological data analysis (TDA) is the emerging framework to process datasets under this perspective. In parallel, topology has proven essential for understanding fundamental questions in physics. Here we report the discovery of topological phase transitions in functional brain networks by merging concepts from TDA, topology, geometry, physics, and network theory. We show that topological phase transitions occur when the Euler entropy has a singularity, which remarkably coincides with the emergence of multidimensional topological holes in the brain network. Our results suggest that a major alteration in the pattern of brain correlations can modify the signature of such transitions, and may point to suboptimal brain functioning. Due to the universal character of phase transitions and noise robustness of TDA, our findings open perspectives towards establishing reliable topological and geometrical biomarkers of individual and group differences in functional brain network organization.

## I. INTRODUCTION

Topology aims to describe global properties of a system that are preserved under continuous deformations and are independent of specific coordinates, while differential geometry is usually associated with the system’s local features [1]. As they relate to the fundamental understanding of how the world around us is intrinsically structured, topology and differential geometry have had a great impact on physics [2, 3], materials science [4], biology [5], and complex systems [6], to name a few. Importantly, topology has provided [7-11] strong arguments towards associating phase transitions with major topological changes in the configuration space of some model systems in theoretical physics. More recently, topology has also started to play a relevant role in describing global properties of real world, data-driven systems [12]. This emergent research field is called topological data analysis (TDA) [13, 14].

In parallel to these conceptual and theoretical advances, the topology of complex networks and their dynamics have become an important field in their own right [15, 16]. The diversity of such networks ranges from the internet to climate dynamics, genomic, brain and social networks [15, 16]. Many of these networks are based upon intrinsic correlations or similarities relations among their constituent parts. For instance, functional brain networks are often constructed by quantifying correlations between time series of activity recorded from different brain regions in an atlas spanning the entire brain [16].

Here we report the discovery of topological phase transitions in functional brain networks. We merge concepts of TDA, topology, geometry, physics, and high-dimensional network theory to describe the topological evolution of complex networks, such as the brain networks, as function of their intrinsic correlation level. We consider that the complex network is related [13, 17] to a multidimensional structure, called simplicial complex, and that a topological invariant (the Euler characteristic, see below) suffices to characterize the sequence of topological phase transitions in the complex network.

The simplicial complex associated with a network is constituted by its set of nodes, edges, triangles, tetrahedrons and higher-dimensional counterparts [13, 17]. A simplicial complex is therefore a multidimensional structure that can be related, e.g., to a functional brain network. We establish the bridge between the above interdisciplinary formalisms by scanning correlation levels in functional brain networks just like one slices the energy levels in a physical system [7, 10, 18–20], or the height function in Morse theory [21] and computational topology [13, 17]. We display in Table 1 and Fig. 1 the proposed analogy between topological changes in a simplicial complex of a brain network and an equipotential energy surface of a physical system, whose structure is determined by its Hamiltonian (energy) function. In a physical system these topological changes occur as energy is increased, while in a functional brain network they emerge as the correlation threshold (or any other similarity measure) is varied. This puts the network’s simplicial complex on a similar footing to the configuration space of the Hamiltonian system’s dynamics, except for the very relevant gap that the underlying Hamiltonian function of actual complex systems is usually unknown and often inaccessible [22]. Under the intrinsic topological approach we propose here, the possibility emerges that this gap can be circumvented (at least partially), thus allowing for significant progress in network theory even in the absence of a Hamiltonian description of the study system.

**TABLE I:** Outline of the analogy between statistical physics and topological data analysis (TDA) of complex networks. Some concepts from statistical physics present analogous topological counterparts, as discussed in the text. Such analogy can be built by merging elements from TDA, physics, topology, differential geometry, and high-dimensional network theory to describe the topological evolution of networks that arise from intrinsic correlations in complex systems, such as functional brain networks.

| Statistical physics *vs.* topological data analysis of complex networks |  |
| --- | --- |
| Interactions based on a Hamiltonian function | Interactions based on similarity measures |
| From microscopic to macroscopic | From local to global |
| Energy, temperature, etc. | Correlation threshold, probability, etc. |
| Boltzmann entropy | Euler entropy |
| Level sets | Filtration process |
| Configuration space | Simplicial complex |
| Phases structure | Betti numbers |
| Usually extrinsic | Usually intrinsic |
| Not necessarily friendly to big data | Frequently friendly to big data |

**FIG. 1:**
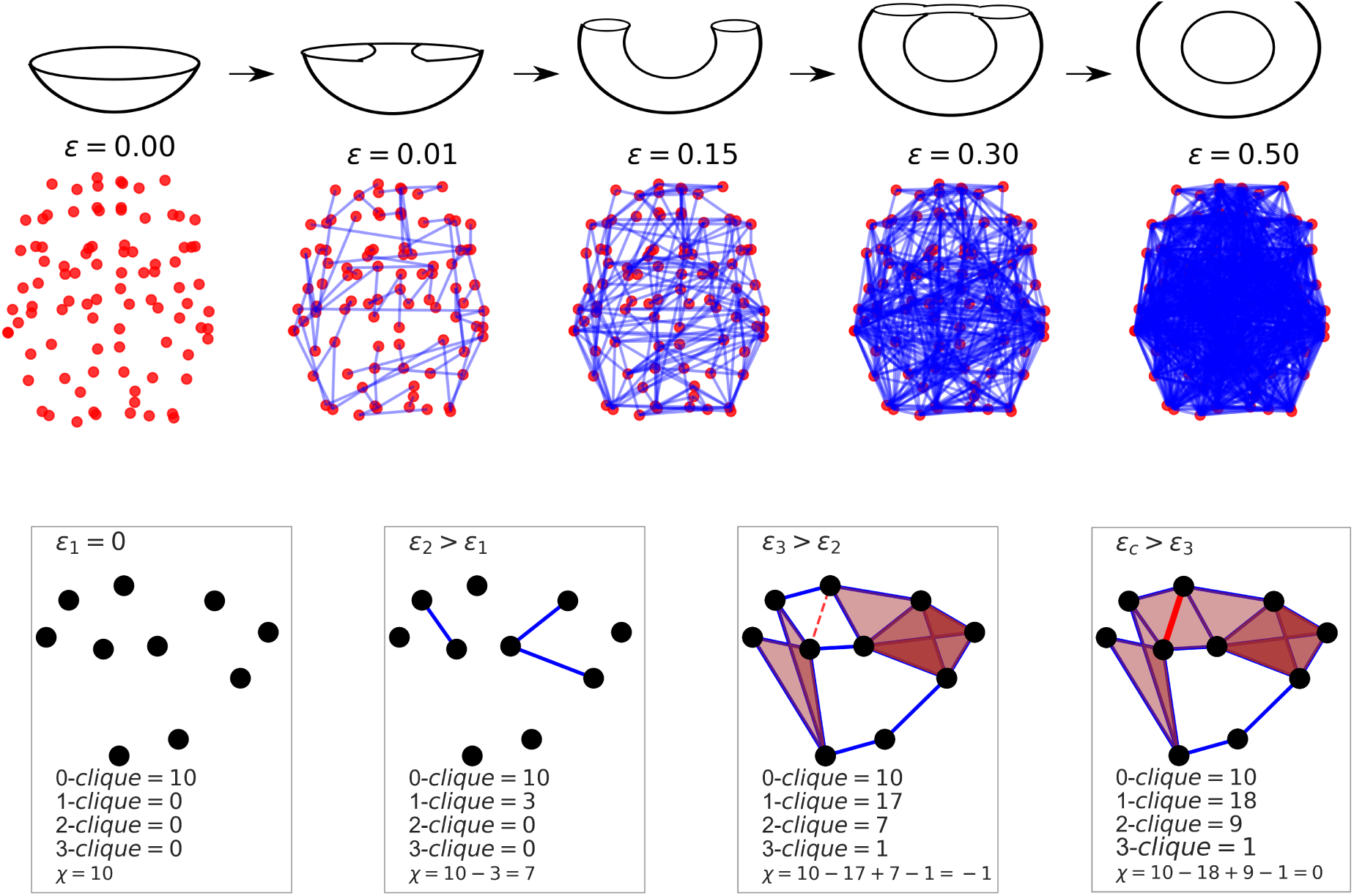
Illustration of the filtration process in a functional brain network and analogy with the level sets of a torus. The middle panel shows regions (red dots) of an individual’s brain from the VUmc dataset. In the filtration process, a functional brain network is built for each value of the correlation threshold *ε* ∊ [0, 1] by assigning an edge linking two brain regions if their normalized correlation level is larger than 1 − *ε*. Therefore, a brain network with no links corresponds to *ε* = 0, whereas a fully connected structure arises for *ε* = 1. As *ε* is enhanced, new edges are gradually attached thus changing the topology of the brain network, which becomes increasingly denser and harder to analyze. This process is analogous to the evolution of level sets in a surface, illustrated as a torus in the top panel, with the increasing of the height parameter level. In this analogy, a topological change takes place in the surface when the level set reaches up a value that delimits cross-section configurations with one and two circles. Topological invariants are able to track those changes in the evolution of both surfaces and networks. Each network has an associated topological structure called simplicial complex, constituted by its nodes (*k* = 1), edges (*k* = 2), triangles (*k* = 3), tetrahedrons (*k* = 4), and higher (*k −* 1)-dimensional counterparts, the so-called *k-*cliques, as shown in the illustrative example of the bottom panel. The alternate sum of the numbers of *k-*cliques determines the Euler characteristic χ. A topological phase transition represents a major change in the network topology, occurring at the value of *ε* for which χ = 0.

We apply our theoretical framework to Erdős-Rényi graphs [23] as well as to brain networks built from resting-state functional magnetic resonance imaging (rs-fMRI) connectivity data publicly available through the Human Connectome Project (HCP) [24, 25] and from VU University Medical Center (VUmc, Amsterdam) [26]. In Erdős-Rényi graphs, the associated random network with uncorrelated links between nodes is exactly solvable [27] and the behavior of the Euler characteristic can be analytically understood. Therefore, Erdős-Rényi graphs can serve as a reliable test ground system in our approach to brain networks. In both cases we find a sequence pattern of topological phase transitions as a function of the linking probability in Erdős-Rényi graphs or the magnitude of intrinsic correlations from rs-fMRI measurements in functional brain networks. It is important to stress, however, that the brain networks are shown here to present a remarkable local structure which is not displayed by Erdős-Rényi random networks.

The Euler characteristic is a central quantity that we address here [17]. Its history goes back to Plato. Consider, for instance, a tetrahedron in three dimensions. It has *N* = 4 nodes, *E* = 6 edges and *F* = 4 faces. Its Euler characteristic is χ = *N* − *E* + *F* = 2. Exactly the same result for this alternate sum is obtained for a cube, an icosahedron, and any convex polyedron. In this sense, they are all homeomorphous to a sphere (a torus, however, yields a different result for χ due to the central hole). Thus, the Euler characteristic is an intrinsic property, which means that it does not depend on the particular parametrization of the given object (see Table 1). The distinction between intrinsic and extrinsic properties of a system was introduced by Gauss [28] and made popular by Nash [29]. The Euler characteristic is also a topological invariant [17]. In the case of surfaces, this means that any two surfaces that can be transformed into each other by continuous deformations share the same Euler characteristic. For complex networks this concept is less intuitive, though one can say that two networks are topologically equivalent if a mapping between their representative graphs preserves their structure, and consequently yields the same Euler characteristic. The logarithm density of the Euler characteristic, introduced as a potential link between thermodynamic entropy and topology [18], was recently defined as the Euler entropy [10]. It presents non-analytical behavior (see below) at points coinciding with the thermodynamic phase transitions in some exactly solvable physical models [7, 18–20], and is arguably useful here as well to set the topological phase transitions that take place in functional brain networks. Moreover, the Euler characteristic is also related to percolation transitions [30], which gives substantial theoretical support and motivation to our approach.

The robustness of our results is corroborated by the computation of another set of topological invariants called Betti numbers [17], which are associated with the number of multidimensional topological holes in the network and therefore also with the brain connectivity [31]. Our work also finds support in recent rigorous results concerning the distribution of Betti numbers in random simplicial complexes [32].

We remark that the TDA framework is able to spatially locate the brain regions that participate most in the connected structures of the functional brain network [33]. With this information in hand, and based on a discrete version of a theorem relating differential geometry to topology [34, 35], here we also provide a possible link between global (topological) and local (geometrical) properties of the brain network. This finding opens the perspective to advance in the quest for a link between local network structure and global functioning in brain networks.

The TDA framework is also particularly valuable since on the one hand the numerical analysis is rather robust against measurement noise [36], while it is also quite sensitive to characterize topological features of multidimensional structures, as recently suggested in [31, 33, 37].

Why is this relevant? Because phase transitions have a universal character by their own nature [38, 39]. Therefore, in the same way that something that boils at 100°C at sea level is very likely to be water, the critical points that locate the topological phase transitions in brain networks have potential to be used as topological biomarkers. In this context, a change in the location pattern of the critical points of the topological phase transitions in a brain network may signal a major change in the correlations among brain regions. It can therefore be fundamentally related to suboptimal brain functioning (see, e.g., [40-42]) and, in perspective, may become relevant to clinical neuroscience of neurological and psychiatric disorders [43].

Our topological approach may thus allow not only to discern novel features of brain organization relevant to human behavior, but also offers new avenues that can impact the investigation of individual differences in a non-biased manner, such as in recent studies of task performance [44] and differentiation between schizophrenia patients and healthy controls [45].

## II. RESULTS

The concept of Euler characteristic of a surface was extended by Poincaré [46] to spaces of arbitrary dimensions as follows. A graph is a structure with nodes and edges. A *k-*clique is a subgraph with *k* all-to-all connected nodes. For example, each individual node is a 1-clique, each edge a 2-clique, each triangle a 3-clique, each tetrahedron a 4-clique, and so on. In other words, one can represent a *k-*clique as a (*k* − 1)-dimensional object. In our notation, Poincare’s extension for the Euler characteristic χ of a graph is computed as an alternate sum of its numbers of *k-*cliques (see below).

### A. Topological phase transitions in Erdős-Rényi graphs

Consider a set of *N* nodes (vertices or points). In an Erdős-Rényi graph [23] any two nodes are connected by a linking edge with probability *p*. Moreover, each edge is attached to the graph independently of every other edge. This implies that each node of an Erdős-Rényi graph is connected, on average, with (*N* − 1)*p* other nodes.

A *random network* in which the uncorrelated connections between nodes occur with probability *p* can be represented by an Erdős-Rényi graph [23]. We can thus investigate the evolution of complex random networks (or Erdős-Rényi graphs) for fixed *N* nodes as a function of the probability parameter *p* ∊ [0, 1]. If *p* = 0 no nodes are connected (empty graph), whereas if *p* = 1 all nodes are fully linked to each other (complete graph).

From the discussion above, in an Erdős-Rényi graph it is possible to identify subgraphs containing *k-*cliques with *k* all-to-all connected nodes (the simplicial complex), and then determine its Euler characteristic. The mean value of the Euler characteristic of Erdős-Rényi graphs with *N* nodes and linking probability *p* is exactly given by the alternate sum [27]

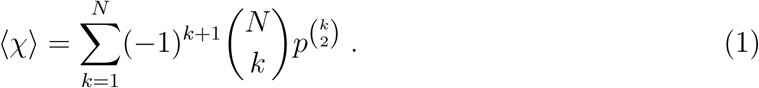

In the above expression, 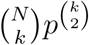 is the mean number of *k-*cliques, since there are 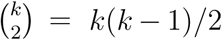 links in a *k-*clique that occur with probability 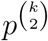. Also, the possible number of choices of *k* nodes from a total of *N* is 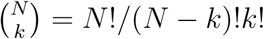.

The Euler entropy [10] of the associated random network is obtained from

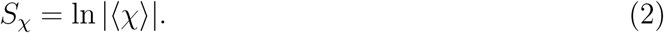

As the linking probability is varied, when 〈χ〉 = 0 for a given value of *p* then the Euler entropy is singular, *S_χ_* → −*∞*. Further, we notice below that a zero of 〈χ〉 and a singularity of *S_χ_* can be related to an important topology change in the Erdős-Rényi graphs. We thus define a topological phase transition in a complex network as the point at which the Euler characteristic is null and the Euler entropy is singular. Actually, this statement finds support in the akin non-analytical behavior of *S_χ_* at the thermodynamic phase transitions of various physical systems [7, 10, 18–20]. In addition, we show below that a distinct set of topological invariants, called Betti numbers, also concurs to verify this assertion independently.

Figure 2(a) displays the Euler entropy of Erdős-Rényi networks with *N* = 50 nodes as a function of the linking probability *p*, calculated from Eq. (1). We first notice the presence of several singularities in *S_χ_* associated with the many zeros of the mean Euler characteristic given by a polynomial of degree 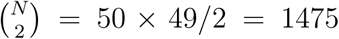, see Eq. (1). The observed sequence of topological phase transitions delimits several phases in the random networks, whose features can be unveiled by the analysis of the Betti numbers.

**FIG. 2:**
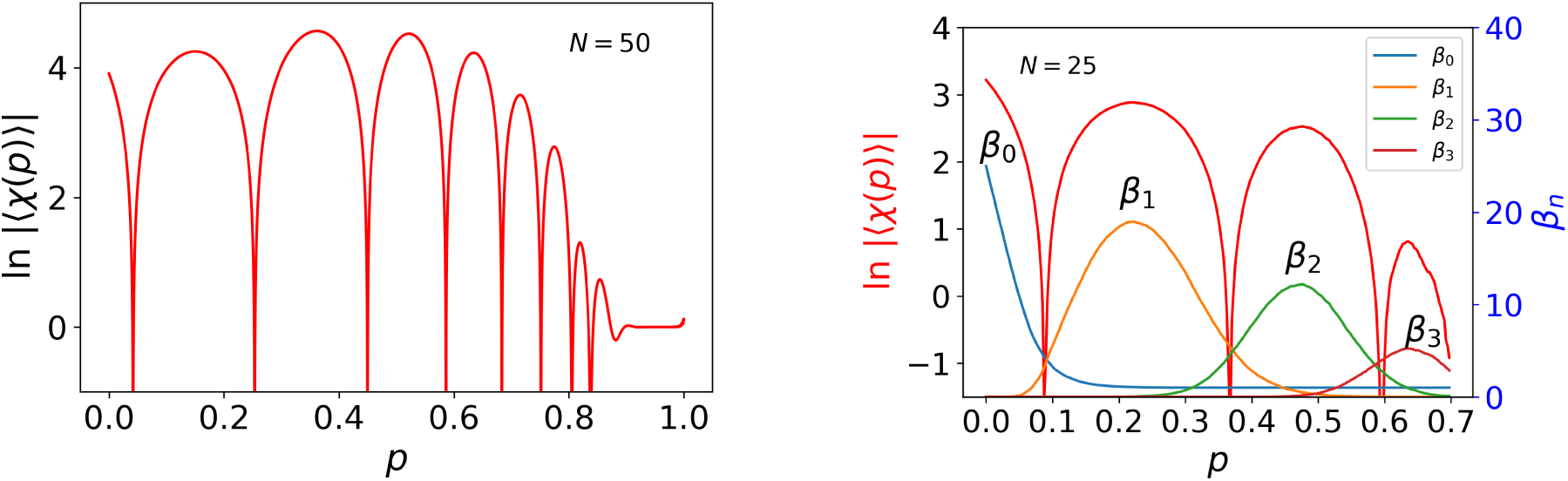
Topological phase transitions in Erdős-Rényi networks. (a) Euler entropy *S_χ_* = ln |〈*χ*〉| as a function of the probability *p* of connecting two nodes in an Erdős-Rényi graph with *N* = 50 nodes. *S_χ_* was determined by Eq. (2), with the average Euler characteristic 〈*χ*〉 exactly given [27] by Eq. (1). The zeros of 〈*χ*〉 or the singularities *S_χ_* → −∞ locate the topological phase transitions in random networks. On the other hand, the topological phases are characterized by the Betti numbers *β_n_*, shown in (b) for *N* = 25. At each given phase, one specific Betti number prevails, featuring which sort of *n*-dimensional topological hole predominates in the network structure. For example, for *N* = 25 and 0.09 ≲ *p* ≲ 0.36 (between the first and second transitions in (b)) we note that *β*_1_ (number of loops or cycles) is much larger than all other *β_n_*’s. The phase boundaries delimited by the dominant *β_n_*’s nicely agree with the loci of the topological phase transitions set by *S_χ_*. As *p* increases, we find a sequence of transitions in which the dominant *β_n_* varies (*n* is added by one unity) every time a transition point is crossed, thus signaling an important change in the topology of the Erdős-Rényi random network.

The Betti number *β*_n_ counts the number of *n*-dimensional topological holes in the simplicial complex of a network [17]. For example, *β*_0_ counts the number of connected components in the network or connected subgraphs in a graph, *β*_1_ is the number of loops or cycles, *β*_2_ represents the number of voids or cavities (like the one in the torus), and so on. We remark that the Euler characteristic can be also expressed as an alternate sum of Betti numbers [17]. In Fig. 2(b) we show the numerical results for *β*_n_ in random networks with *N* = 25, indicating that the topological phase which sets in between the *n*-th and (*n* + 1)-th transitions corresponds very closely to the range in *p* where the Betti number *β_n_* prevails. E.g., for *N* = 25 and 0.09 ≲ *p* ≲ 0.36 (between the first and second transitions) we note that *β*_1_ is much larger than all other *β_n_*’s. This means that loops (*n* =1) are abundant in such networks in this range of *p*. As *p* increases, we find a sequence of dominant Betti numbers *β_n_*, starting from *n* = 0, that change (i.e., *n* is added by one unity) every time a topological phase transition is crossed. In other words, while the location of the transitions is determined by the singularities of the Euler entropy *S_χ_*, the Betti numbers *β_n_* characterize which kind of multidimensional hole prevails in each topological phase.

Another interesting connection can be set with percolation theory. Indeed, the first transition in the sequence shown in Fig. 2(b) takes place at a value *p* = *p_c_* that nearly corresponds [23, 32] to that of the percolation phase transition, which gives rise to the giant connected component of the Erdős-Rényi graph containing most of the network nodes. This finding is supported by recent rigorous results [32, 47] from the distribution of Betti numbers that actually span all phase transitions, not only the first one, as well as from the topological data analysis of continuous percolation with disk structures [48]. In fact, the percolation transition in two-dimensional lattices is known [49] to occur in the vicinity of the only nontrivial zero of their Euler characteristic. Since the lattice percolation problem involves the counting only of nodes, edges and faces in the Euler characteristic [30, 49], our results can further contribute to the detection of percolation transitions in higher-dimensional objects (triangles, tetrahedrons, etc.), a concept that was actually defined only very recently for complex systems [50]. Indeed, for systems where the distribution of the Betti numbers are concentrated in an interval in the limit *N* ≫ 1, the percolation of *k*-cliques (or *k*-simplices) might occur in the vicinity of the critical point in which *β_k_* ≈ *β_k_*_+1_ [32, 47]. Therefore, in the thermodynamic limit and under conditions analogous to those of [32, 47], the zeros of the Euler characteristic suffice to detect percolation transitions in higher-dimensional objects of the associated simplicial complex. We comment that this result deserves further investigation.

We can also comment on the order of the topological phase transitions in these complex networks. It is known [32] that the percolation transition in Erdős-Rényi random graphs is of second order [38, 39] in the sense that, e.g., the relative number of nodes linked by edges (2-cliques) in the giant component shifts without discontinuity from zero for *p* < *p_c_* to a non-null value for *p > p_c_*. Nevertheless, the order of the subsequent transitions in Fig. 2(b) deserves further investigation. On the one hand, the similar logarithmic singular behavior of *S_χ_* could suggest that the subsequent transitions are of second order as well. Notwithstanding, one cannot also discard a discontinuous first order transition as observed in [32], in which the concept of giant component was extended to higher-dimensional *k*-cliques structures, such as triangles (*k* = 3), tetrahedrons (*k* = 4), etc. Thus, a numerical study on the connectivity of (*k >* 2)-cliques and comparison with [32] may shed light on this point. Analogous arguments might be also applied to brain networks, see below.

### B. Topological phase transitions in functional brain networks

Most real complex networks are not random networks or Erdős-Rényi graphs [15]. In particular, the connections between any two nodes (brain regions or voxels) in a functional brain network are not randomly established with probability *p*, but are instead determined from the magnitude of their intrinsic correlations. One may thus ask if topological phase transitions also occur in actual data-driven complex networks, as functional brain networks.

To explore the relevance of the TDA framework in neuroscience, we analyze two different rs-fMRI datasets to construct functional brain networks. On the one hand, raw correlation data between *N* = 177 brain regions of 986 individuals were considered from the Human Connectome Project (HCP dataset) [24, 25], while, on the other hand, z-score values of correlations among *N* = 92 brain regions of 15 healthy individuals from VU University Medical Center (VUmc dataset) were investigated [26].

In the brain networks considered here, each node corresponds to a different region of the brain. Each individual is associated with a *N* × *N* matrix *C_ij_ o*f correlations between regions *i* and *j*, which is built from the time series obtained from the rs-fMRI measurements (see Methods for details). The filtration process in the TDA analysis involves the definition of correlation threshold *ε ∈* [0, 1] so that, for the brain network of a given individual, two regions *i* and *j* are connected by an edge if the associated matrix element *C_ij_* is such that |*C_ij_*| *>* 1 − *ε*. This implies that for *ε* = 1 all regions in the brain are connected, while there are no connections if *ε* = 0. It is thus possible to follow the evolution of the functional brain network of each individual as a function of the correlation threshold level *ε*, in analogy to the previous analysis with *p* of the Erdős-Rényi random networks.

The Euler characteristic of the functional brain network of each individual is not given by Eq. (1), as in the case of random networks, but is expressed by the alternate sum of the numbers *Cl_k_* of *k*-cliques present in the simplicial complex of the network for a given value of correlation threshold *ε* [17],

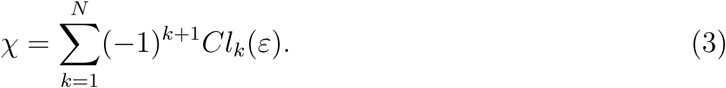

The Euler entropy *S_χ_* is thus obtained from *χ* as in Eq. (2).

Figure 3(a) shows the Euler entropy as function of *ε* for the HCP dataset. As discussed in the Methods section, the numerical computation of the numbers of *k*-cliques in brain networks becomes an increasingly (exponentially) difficult task as *ε* raises and more edges are progressively attached. This is actually an NP-complete problem, however without a closed analytical expression available for the Euler characteristic of the brain network, such as Eq. (1) for the Erdős-Rényi graphs. This is in fact the reason why we did not go beyond *ε* =0.60 with steps Δ*ε* = 10^−2^ in Fig. 3(a), even analyzing close to this upper limit data from only 420 out of the 986 individuals in the HCP dataset. Nevertheless, the Euler entropy averaged over the individuals (blue line) clearly shows the presence of three singularities in this range of *ε*, which correspond to the loci of the topological phase transitions taking place in these functional brain networks.

**FIG. 3:**
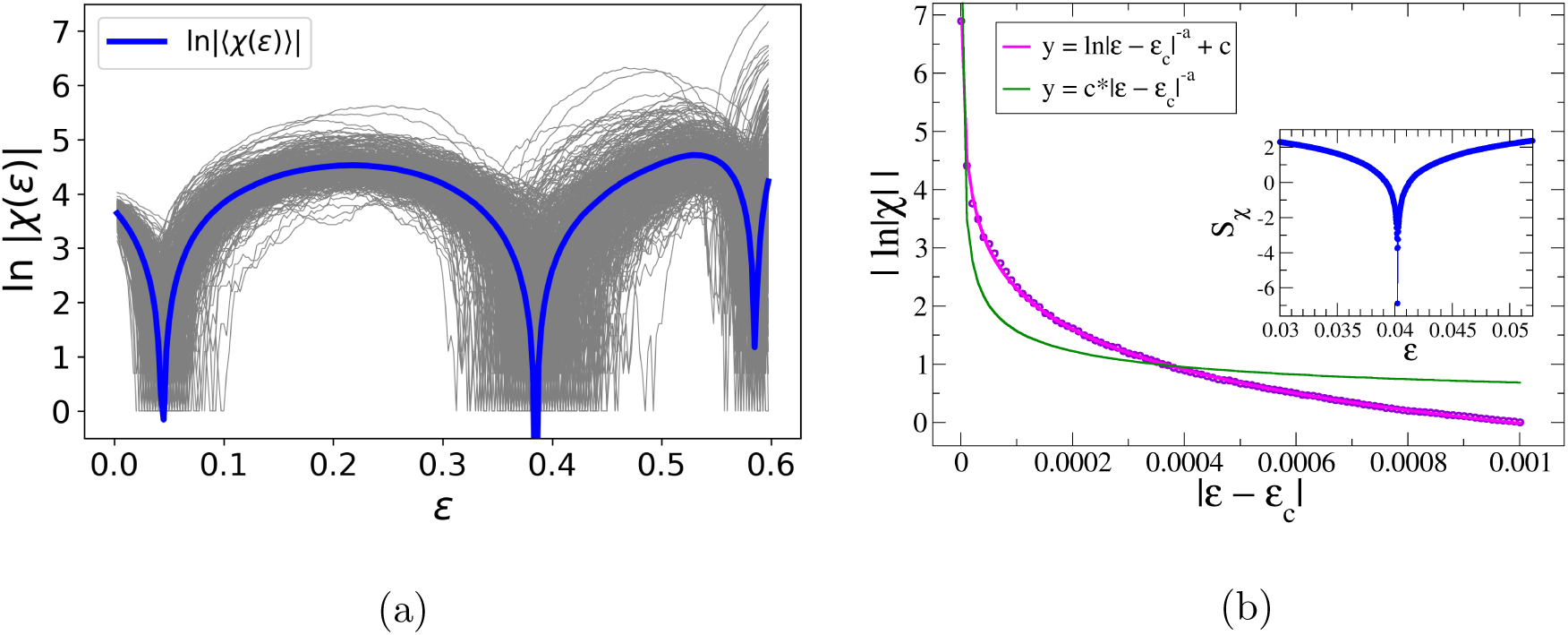
Topological phase transitions in functional brain networks. (a) Euler entropy *S_χ_* = ln |*χ*| as a function of the correlation threshold level *ε* of functional brain networks from the HCP dataset. Each gray line represents an individual’s brain network, whereas the blue line depicts their average. A sequence of three topological phase transitions in the brain networks is identified in this range of *ε* at the deepening points of *S_χ_* (blue line). (b) A quite detailed analysis of *S_χ_* near the first transition at *ε_c_* ≈ 0.04 was performed using steps Δ*ε* = 10^−5^ (inset), in contrast with Δ*ε* = 10^−2^ used in (a). The excellent fit (magenta line) to the data (blue circles) confirms the logarithmic singularity *S_χ_* = ln |*ε* − *ε_c_*|^α^ + *c*, with *c* as a constant and best-fit exponent *α* =1.004 (*α* =0.985) as the transition is approached from below (above), in nice agreement with the theoretical prediction *α* = 1. We also include for comparison the unsuccessful fit (green line) to the alternative form *S_χ_* = *c*|*ε* − *ε_c_*|^α^.

We comment that these singularities of *S_χ_* resemble the singular cusp behavior of the Euler entropy reported at the phase transition of some physical systems [7, 10]. Indeed, this finding is corroborated in Fig. 3(b) by the excellent fit of the data to the expression *S_χ_* = ln |*ε* − *ε_c_*|*^α^* + *c*, where *c* is a constant, as *ε* approaches the first transition point at *ε_c_* ≈ 0.04 with much narrower steps Δ*ε* = 10^−5^ for the whole HCP dataset of 986 individuals. The inset of Fig. 3(b) shows the detailed behavior of *S_χ_* in the critical region very close to *ε_c_* [38, 39]. The best-fit value of the exponent is *α* =1.004 (*α* =0.985) as the transition is approached from below (above). These results nicely agree with the prediction *α* = 1 from the Euler characteristic written as a polynomial function with set of zeros {*ε_c,i_*}, that is, *χ* = ∏*_i_*(*ε* − *ε_c,i_*). By contrast, we also include in Fig. 3(b) the unsuccessful attempt to fit the alternative form *S_χ_* = *c*|*ε* − *ε_c_*|*^α^*.

In order to provide further support to these results and additional characterization of the topological phases in the brain networks, we also calculate the Betti numbers *β_n_*. Figure 4 presents results from both HCP (top panel) and VUmc (bottom panel) datasets. In the former case we considered data from a subset of 712 individuals, which limited our analysis up to *ε* =0.50.

**FIG. 4:**
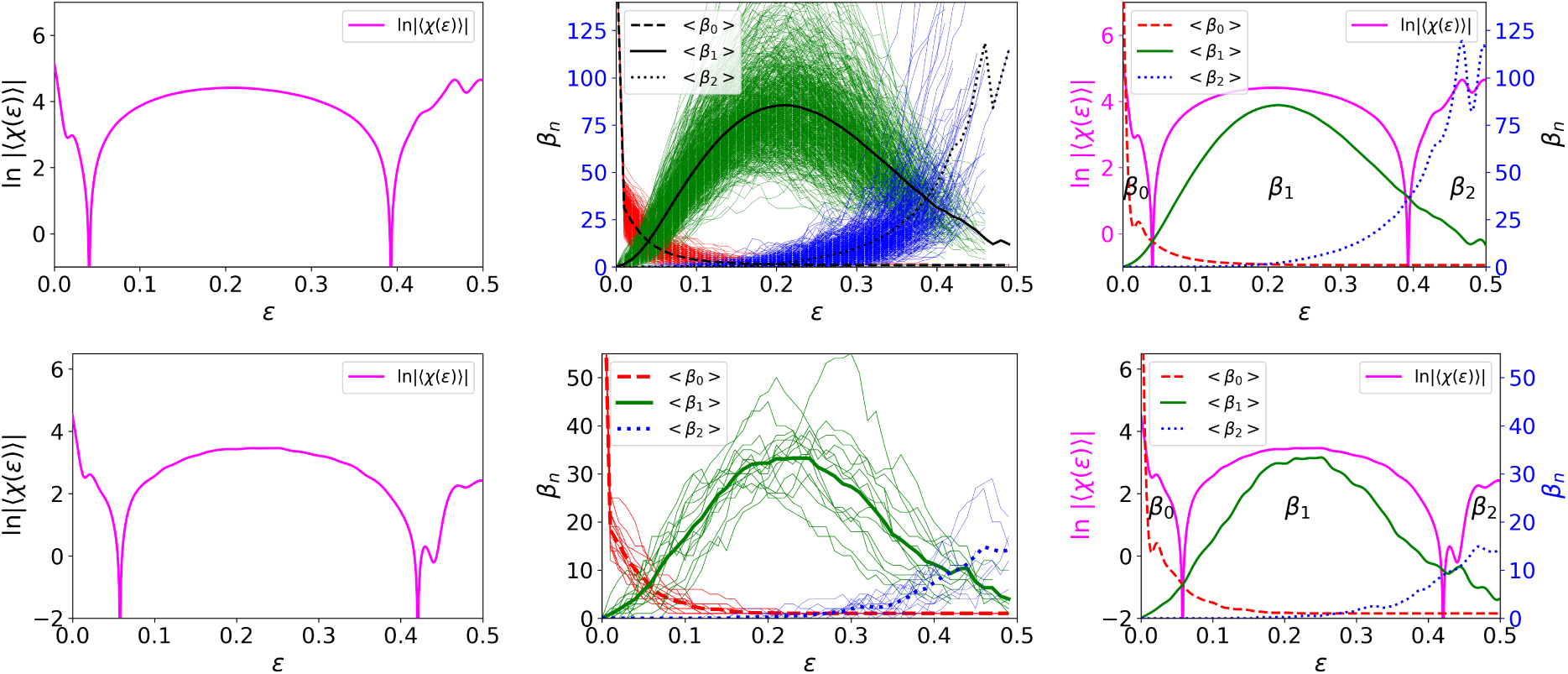
Characterization of topological phases in functional brain networks from two different datasets. Euler entropy (left plots) and the three first Betti numbers (central plots) as a function of the correlation threshold *ε* of functional brain networks from the HCP (top panel) and VUmc (bottom panel) datasets. Each thin line in the central plots represents an individual’s brain network, whereas thick lines depict their averages. As in Erdős-Rényi random networks, see Fig. 2, the right plots show a fine agreement between the phase boundaries delimited by the dominant Betti numbers and the loci of the topological phase transitions set by *S_χ_*. For values of *ε* below the first transition, where *β*_0_ prevails, the brain network is characterized by a high fragmentation level, with the number of connected components larger than that of cycles, voids, etc. Between the first and second transitions, an important topological change takes place in the brain networks and the number of loops or cycles overcomes the number of connected components. After the second transition and up to the maximum threshold reached in our numerical analysis, *β*_2_ becomes the dominant Betti number, indicating the proliferation of voids or cavities in the densely correlated functional brain networks, along with the vanishing of cycles.

The left and central plots of Fig. 4 show, respectively, the Euler entropy and the first three Betti numbers, *n* =0, 1, 2. As evidenced in their average values displayed together in the right plots, in both datasets we notice a remarkable agreement between the transition points, determined by the singularities of *S_χ_*, and the boundaries of the topological phases characterized by each dominant *β_n_*, as observed theoretically in [32]. For instance, in the HCP dataset the range between the first (*ε* ≈ 0.04) and second (*ε* ≈ 0.39) topological transitions nicely coincides with the phase in which *β*_1_ is larger than both *β*_0_ and *β*_2_. In this regime, the brain network is plenty of one-dimensional topological holes (loops). In contrast, in the topological phase for *ε* ≳ 0.39 and up to the maximum *ε* reached in our study the dominant *β*_2_ indicates the proliferation of voids or cavities in the more densely correlated network. Furthermore, the large value of *β*_0_ below the first phase transition at *ε* ≈ 0.04 implies a topological phase with substantial fractioning of the brain network into several disconnected components. In this sense, the first transition in Fig. 3(a) resembles the first percolation transition in Erdős-Rényi random networks discussed above, below which the giant connected component cannot establish.

### C. Connection between local network structure and brain function: from geometry to topology

A fundamental question that emerges in the present scenario concerns the relation between the global (topological) features and local (geometric) properties of the brain network. Moreover, another rather important related issue is the understanding of the connection between the network structure and brain function.

A key starting point in this direction might be the study of the individual nodes and their contributions to form all-to-all connected *k*-cliques structures (triangles, tetrahedrons, etc.) that constitute the simplicial complex of the brain network. Indeed, the potential role of *k*-cliques for the link between structure and function has been recently addressed in the Blue Brain Project [51], where it was observed that groups of neurons are arranged in directed *k*-cliques that give rise to topological structures in up to eleven dimensions (i.e., maximum *k* = 12, see below) emerging in neocortex tissues.

In this context, a suitable local quantity to analyze is the participation rank of each node (brain region) in the k-cliques [33]. It is denoted by Cl*_ik_*(*ε*) and counts the number of *k*-cliques in which node *i* participates for a given *ε*. Here we track the node participation rank in the brain networks as a function of *ε*. Our approach thus differs from that of Ref. [33], in which a fixed correlation threshold was considered.

We start by illustrating in Fig. 5 the emergence, as *ε* increases, of 3-cliques only (triangles) in the functional brain network of an individual from the VUmc dataset. We do not show results for values higher than *ε* =0.15 because in this case the number of triangles enhances considerably, hampering the visualization (see, e.g., the distribution of links in Fig. 1 for *ε >* 0.15). The color bars indicate the percentage of 3-cliques in which each node takes part. Remarkably, we notice that the distribution of the participation ranks in cliques is not homogeneous over the nodes, indicating a spontaneous differentiation among nodes as the correlation threshold raises. An html version of this figure is available in the Supplementary Information, in which it is possible to rotate the brain and identify the locations of the regions that participate most in the 3-cliques structures.

**FIG. 5:**
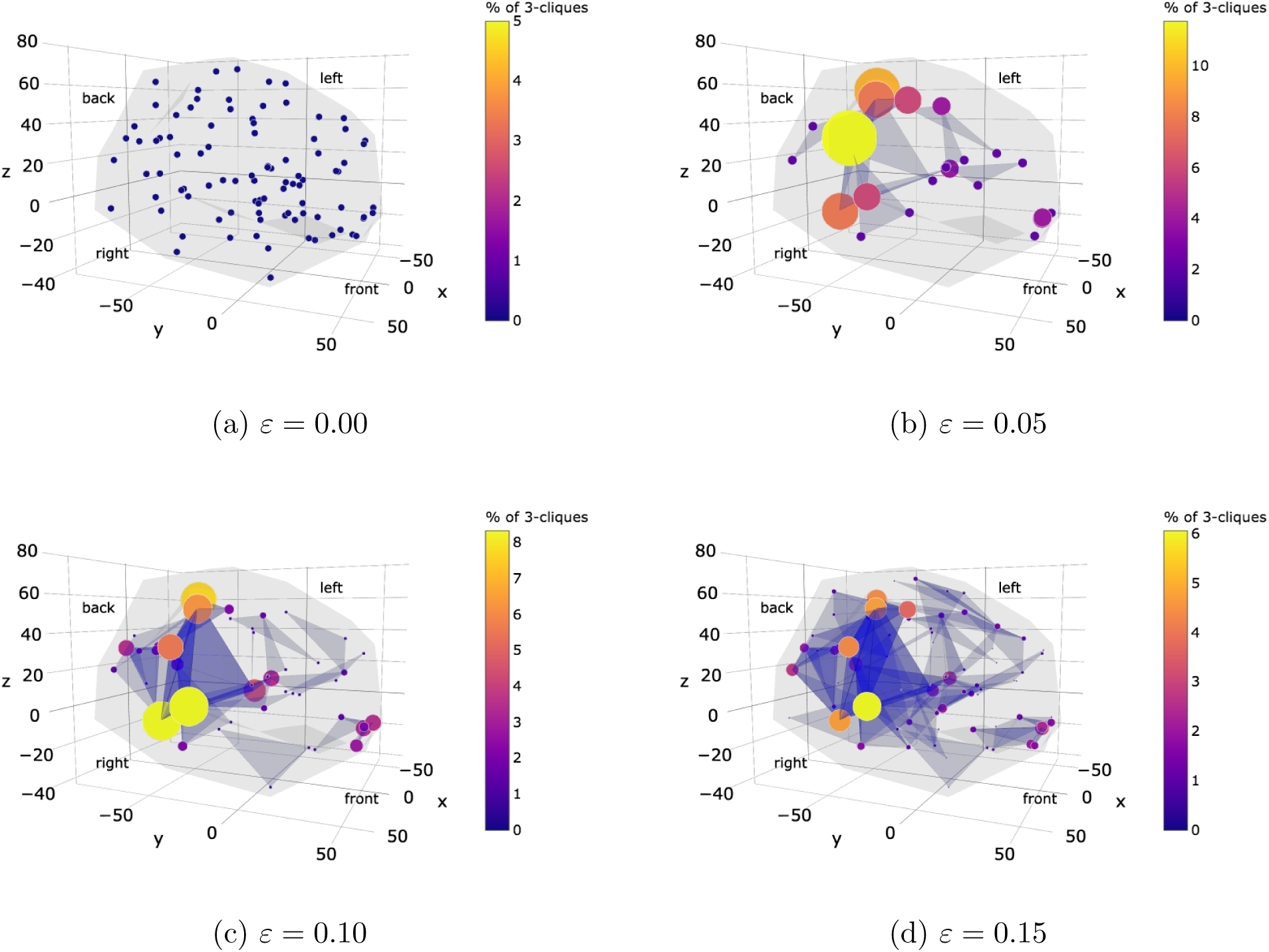
Participation of brain regions in triangular-connected structures of a functional brain network. Evolution with the correlation threshold *ε* of the normalized participation rank of each local brain region (small blue dots in (a)) in 3-cliques (triangles) structures in the brain network of an individual from the VUmc dataset. As *ε* increases, the color code and size of nodes, which are proportional to the participation in 3-cliques, indicate that an spontaneous differentiation of the role of distinct brain regions emerges in the functional brain network.

Figure 6 depicts the node participation rank in all *k*-cliques (we do not draw the triangles, tetrahedrons, etc., for clarity; see also the Supplementary Information). We notice that the nodes which participate in the largest numbers of *k*-cliques are ranked as the most important ones (according to the clique structure) [33]. We also observe that nodes with high participation seem to be similar to hubs defined on the basis of more conventional network measures [15, 33], a point that can be explored in further studies, see e.g. [52]. Since *k*-cliques are essential to compute topological quantities, such as the Euler characteristic and Betti numbers, the access to the distribution of node participation in *k*-cliques can contribute to the understanding, from a local perspective, of the topological phases and topological phase transitions in the brain network. Moreover, the identification of the spatial location of the main cycles and cavities in the brain can be also relevant in the quest for a link between local structure and brain function, with potential applications to the local characterization of functional brain networks [33].

**FIG. 6:**
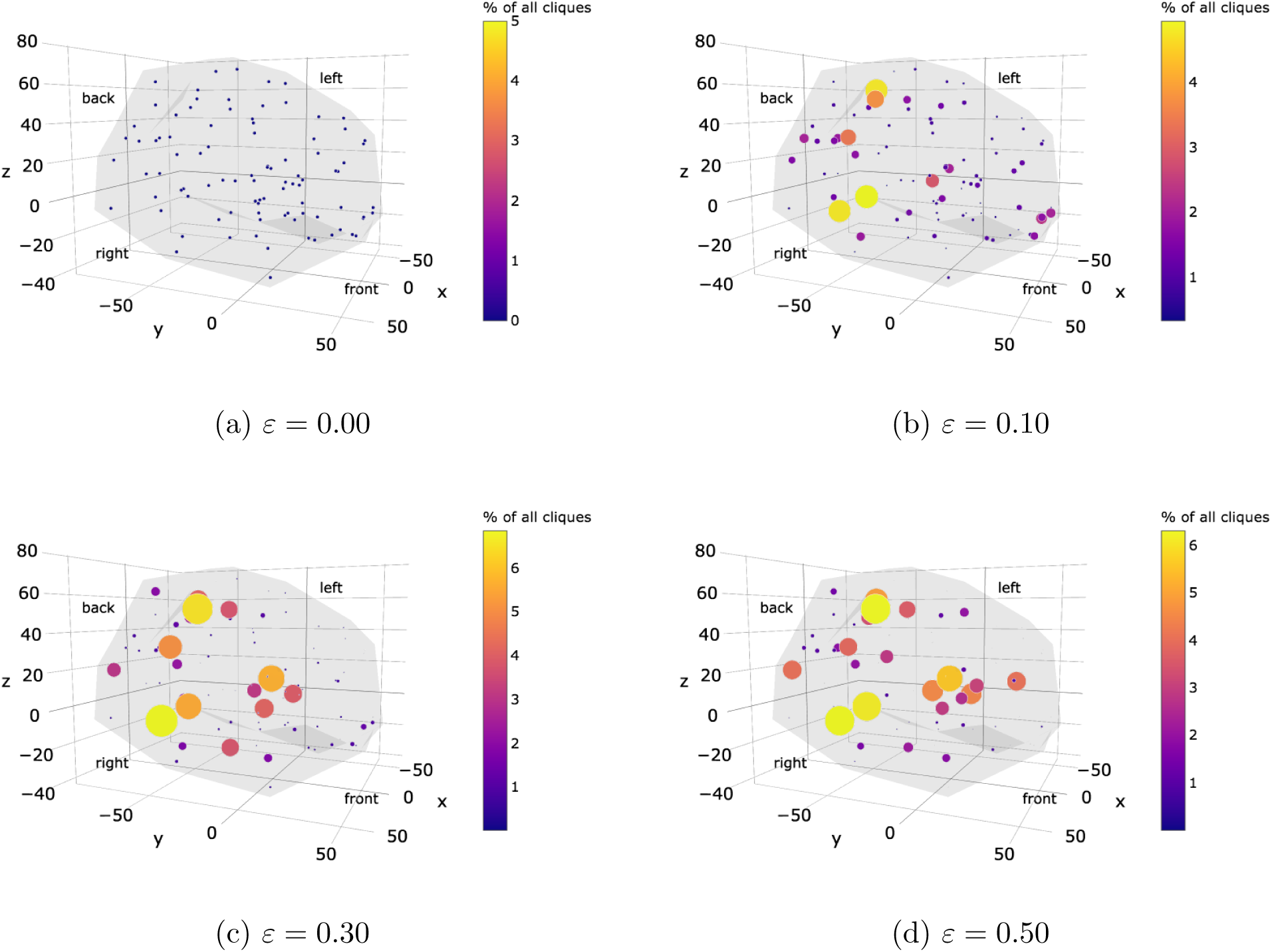
Participation of brain regions in all connected structures of a functional brain network. The same study of Fig. 5 but for all connected (*k*-cliques) structures in the functional brain network. The most important brain regions (regarding the k-clique structure) are the ones with higher participation ranks. The distribution of participation ranks in all *k*-cliques allows to compute the curvature of the brain network at each brain region, thus providing a local-to-global (geometry-topology) connection (see Fig. 7). Moreover, the spatial analysis of participation ranks also opens the perspective for establishing a link between local network structure and brain function [33].

The connection between differential geometry and topology is also useful in this context. In differential geometry it is possible to define the concept of curvature at a given point of a surface essentially as a local measure of how the surface bends in distinct directions [1, 28]. This concept can be extended to complex networks so that one possible definition of the network curvature *κ_i_* at node *i* in terms of the node participation rank is given by [35]

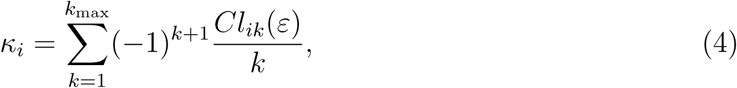

where *Cl_i_*1 = 1 since each node *i* participates in only one 1-clique (the node itself). Also, *k*_max_ represents the maximum number of nodes that are all-to-all connected in the network. Thus, since *k* all-to-all connected nodes form a (*k* − 1)-dimensional object, as discussed above, then one can say that the simplicial complex of the network comprises topological structures in up to *k*_max_ − 1 dimensions. We also observe that Eq. (4) has been successfully applied to complex systems in up to two dimensions [53].

A possible route to connect the geometry (local curvature) of a continuous surface to its topology (Euler characteristic) is given by the Gauss-Bonnet theorem [28]. In a simplicial complex, a discrete version of the theorem can be expressed as [35]

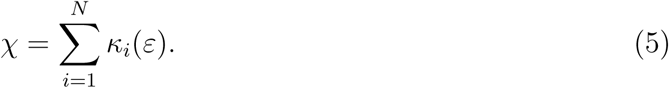

This connection has been explored in complex systems [53]. We also comment that the introduction of the node participation rank in [33] aimed to study the distribution of *k*-cliques in the human connectome, thus relating the appearance of cycles and cavities to the local network structure. Here, however, we intend to understand the topological phase transitions in brain networks from a local perspective, and to this end we compute the network curvature at each node, which also requires the knowledge of the node participation rank, see Eqs. (4) and (5).

In Fig. 7 we display results for the distribution of curvatures *κ_i_* at the nodes (brain regions) of the functional brain network of the same individual of Figs. 5 and 6. We choose three values of *ε*: one between the first and second topological transitions (*ε* =0.200), one right at the second transition (*ε* =0.372 ≈ *ε_c_*), and one after the second transition (*ε* =0.500). (Note from the bottom panel of Fig. 4(b) that the threshold value at the second transition averaged over all individuals in the VUmc dataset is slightly higher, *ε_c_* ≈ 0.42.) First, we remarkably verified the Gauss-Bonnet theorem, Eq. (5), for all *ε*’s considered. This attests the self-consistency of our TDA approach as well as the robustness of the local-to-global connection in brain networks.

**FIG. 7:**
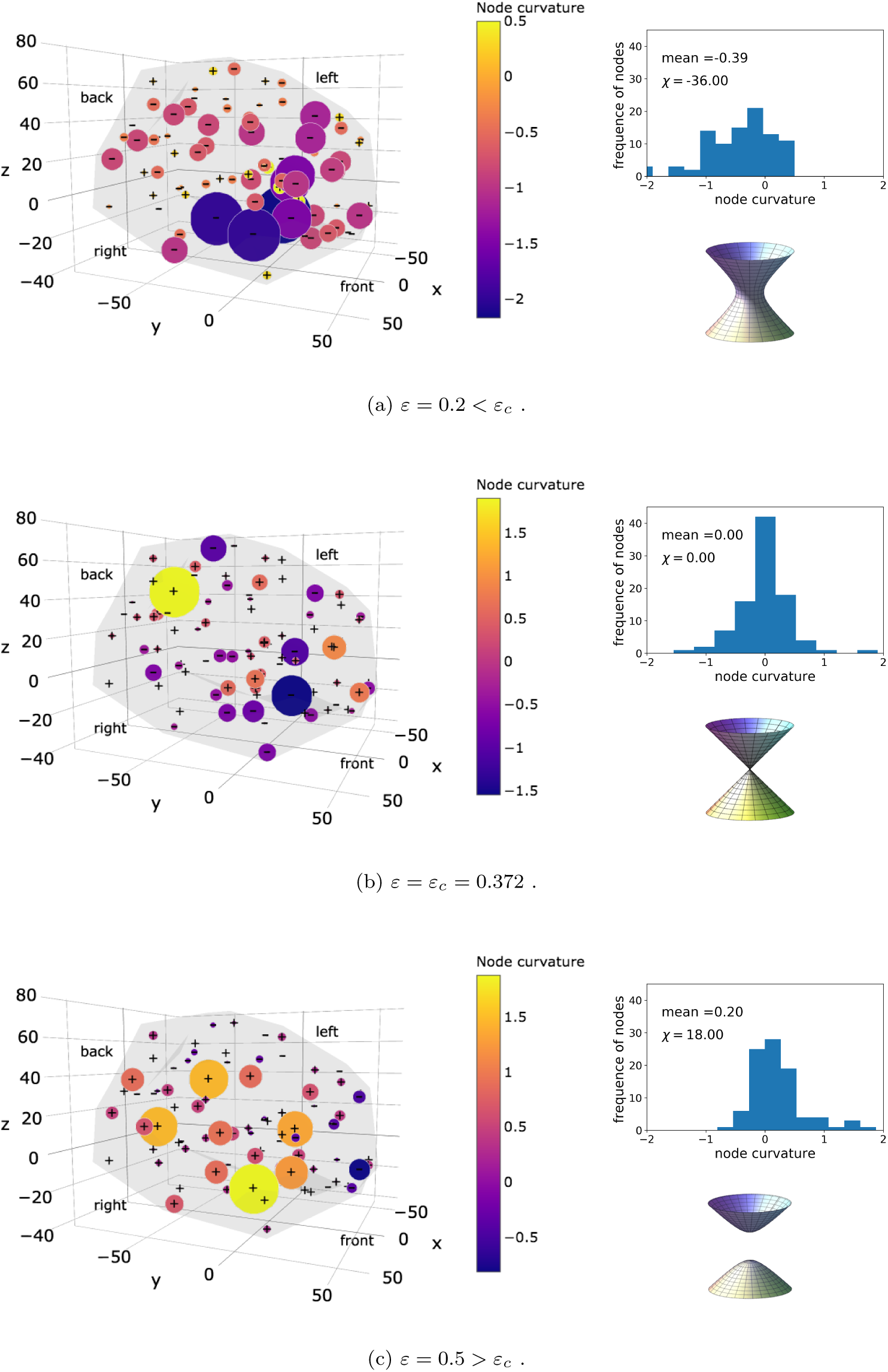
Local-to-global connection in a functional brain network. (a) An one-sheet hyperboloid (top surface) has negative curvature. (b) By tightening the neck of the hyperboloid, it is deformed into a cone (mid surface), which has zero curvature. (c) By detaching the pieces of the cone and smoothing it, one finds a two-sheet hyperboloid (bottom surface), which has positive curvature. A similar evolution occurs for the curvature of the nodes in a functional brain network as a function of the threshold *ε*. The left brain plots illustrate the spatial location of negative, null and positive curvatures, whose distribution is displayed in the histograms. At the topological phase transition (*ε* = *ε_c_*) of the brain network the mean curvature is null (mid panel). This point separates brain networks with negative (*ε < ε_c_*) (top panel) and positive (*ε > ε_c_*) (bottom panel) mean curvatures, just as the cone separates the one-sheet and two-sheet hyperboloids, exactly at the the zeros of the Euler characteristic and singularities of the Euler entropy.

Moreover, we further point out that the local curvature of most nodes at the topological transition is null (locally “flat” network), as also illustrated in the spatial brain representation in Fig. 7(b). Indeed, the mean local curvature is identically zero at the transition point *ε* =0.372 ≈ *ε_c_*. This contrasts with the negative mean curvature (−0.39) before the transition at *ε* =0.200, and the positive mean curvature (0.20) after the transition at *ε* =0.500. Figure 7 also illustrates this picture in the form of a one-sheet hyperboloid surface of negative curvature at *ε* < *ε_c_*, a locally-flat cone of null curvature at *ε* = *ε_c_*, and a two-sheet hyperboloid with positive curvature at *ε* > *ε_c_*. Remarkably, from Eq. (5) a null mean curvature implies *χ* = 0, which is also consistent with the assertion of the zeros of the Euler characteristic and singularities of the Euler entropy to set the location of the topological phase transitions in functional brain networks (Section 1.2).

The local network structure defined both by the node participation rank in *k*-cliques and local curvature allows to easily differentiate functional brain networks from Erdős-Rényi random networks. Indeed, in Erdős-Rényi graphs those quantities are homogeneous over the nodes, which contrasts markedly with Figs. 5-7. In the random network case, due to the same probability *p* of setting an edge linking any pair of nodes, the colors and size of circles in Figs. 5-7 would appear essentially undifferentiated.

Finally, the above scenario resembles considerably the behavior of some physical systems in the vicinity of phase transitions, at which the curvature of the equipotential energy surface in the configuration space is asymptotically null [20, 54]. Indeed, an analogous evolution of the conic and hyperboloid surfaces shown in Fig. 7 can be also found for the equipotencial surface of some magnetic spin systems [20]. In this context, it is important to remark on a very recent empirical description [55] of the Betti numbers associated with different smells that can be described theoretically using a three-dimensional hyperbolic space, which has negative curvature. In fact, the relation between Betti numbers and curvature is well known for surfaces [56], and thus a possible extension of such relation for a simplicial complex deserves further investigation. All these findings reinforce the connection between topology, geometry, and physics to evidence and characterize the topological phase transitions in functional brain networks.

## III. DISCUSSION AND CONCLUSIONS

The discovery that topological phase transitions occur in functional brain networks brings in its wake a number of striking consequences.

The universality principle of phase transitions states that a few properties of the system suffice to determine its macroscopic behavior close to the transition [38, 39]. In this sense, systems that share those properties, but are microscopically distinct, display the same behavior near the transition. Conversely, the critical value of the control parameter at which the phase transition takes place is a system’s ‘fingerprint’ that can be used to differentiate it from others. Here we have located and characterized a sequence of topological phase transitions, associated with important changes in the topology of functional brain networks, by employing the topological data analysis (TDA) approach along with tools and concepts from topology and differential geometry and their relation to theoretical physics.

From the framework point of view, our results give strong support for the use of the Euler characteristic and entropy, Betti numbers, node participation in *k*-cliques structures, and distribution of local curvature as topological and geometrical markers closely associated with functional brain networks in particular and data-driven complex networks with embedded correlations in general. In this sense, our work, together with other recent initiatives that also applied topology ideas to neuroscience [31, 33, 37, 44, 45, 51, 57], reinforces the conjecture by Zeeman in 1965 that topology is a natural mathematical framework to capture the underlying global properties of the brain [58].

The possible connection between the local (geometrical) and global (topological) structures in brain networks has been firmly established by our results. The topological phase transitions are equally determined both from topological quantities (zeros of Euler characteristic and singularities of Euler entropy), as well as from local geometrical measurements, through the null mean curvature at nodes (brain regions) of the functional brain network. From these quantities, we also found a remarkable confirmation of the discrete Gauss-Bonnet theorem in brain networks.

The potential implications of this approach are overarching. On the one hand, the fact that our TDA analysis is multidimensional consequently yields an ampler range of available information on the brain’s intrinsic structure and connections, without necessarily increasing the number of (subjectively chosen) working parameters.

Moreover, the identification of reliable biomarkers of individual and group differences is a crucial issue in clinical neuroscience and personalized medicine. In this context, the location of the topological phase transitions in functional brain networks has potential to be used as topological biomarkers. The TDA framework allows to classify the location sequence of topological transitions in brain networks both at the individual and collective levels, thus enabling direct comparisons with control group standards. A change in the location pattern of the topological transitions may signal an important change in the correlations among brain regions. It can therefore be fundamentally related to possible suboptimal brain functioning [40–42], and may become a relevant precision tool in the clinical diagnosis of neurological and psychiatric disorders [43]. Further studies on this issue will be left for future work.

We have also shown that our approach is able to spatially locate the brain regions that participate most in the connected structures of the functional brain network. This finding may give rise to potential advances in the quest for a proper link between local network structure and brain function.

The joint use of the TDA approach with concepts from topology, geometry, and physics is certainly not restricted to the study of brain networks. Actually, this interdisciplinary framework can be readily applied to look into topological and geometrical properties of other complex networks, such as proteomic, metabolomic, and gene expression, to name a few. These ideas, together with further theoretical insights from TDA, may allow a qualitative change in big data analysis, moving from theory-blind machine learning to a firmer ground based on an intrinsic way of connecting empirical data to the theoretical formalism of these disciplines.

Lastly, this work also relates to one of the most important current scientific debates: Does one miss causality when studying complex phenomena from the big data perspective? Most large datasets generated nowadays under the big data approach are concerned with analyses and predictions based upon statistical correlations only, therefore usually lacking causality relations established from a solid theoretical background [59]. In this sense, putting together ideas from those well-grounded theoretical fields to treat big data-driven complex systems may appear as a promising way to settle this question [22]. Just like the analogy proved to be fruitful here, there are many other results connecting topology, geometry, and theoretical physics (see Methods) that may help one to navigate and improve the vast repertoire of tools available in TDA. In the same way that Riemannian geometry enabled the development of general relativity, the emerging field of TDA has potential to trigger principles that could boost the understanding of the big data revolution from this interdisciplinary perspective.

In conclusion, the discovery of topological phase transitions in brain networks can open the perspective for establishing reliable topological and geometrical biomarkers of individual and group differences. The joint use of interdisciplinary concepts and tools under the TDA approach might change the way complex systems data are analyzed, and can contribute to solve currently open significant questions related to complex networks in various fields, including neuroscience and medicine.

## IV. METHODS

### A. Theoretical support from the connection between topology, geometry, and physics

The impact of the use of tools and ideas from topology and differential geometry has permeated many areas of science. In fact, by applying Riemannian differential geometry, Einstein in his theory of gravity (general relativity) has changed our conception of the space-time structure in the presence of massive cosmological objects [60]. It still thrives and has led to surprising theoretical findings and recent groundbreaking experimental discoveries, including black holes and gravitational waves [61].

On the other hand, topology has played a seminal role in several areas of modern physics. Indeed, topological quantum field theory is nowadays a mature topic [62, 63], while the description of several phenomena in condensed matter physics has achieved a deeper understanding by taking advantage of topological concepts [2, 3]. The bridge between topology and differential geometry can be established through theorems that link local properties of a system with global ones, such as the Gauss-Bonnet theorem [28].

One of the greatest achievements of the use of topology ideas in physics concerns the introduction of the concept of topological phase transition [2, 3]. Examples range from the integer and fractional quantum Hall effects to topological insulators, including the quest for quantum computation [2, 3, 64, 65]. As a consequence, the notion of the topological phase of matter has emerged. Its distinction from the usual phases of matter that undergo equilibrium phase transitions has been emphasized, notably the absence of symmetry breaking or a local order parameter as well as the role played by topological invariants that provide stability (protection) against perturbations [2, 3, 64, 65]. Also, the identification of topological phases, either in quantum or classical physical systems, has recently received special attention in the context of machine learning ideas [66].

In addition, much effort has been also devoted to characterize phase transitions in the configuration space of a physical system from a topological viewpoint [7, 8]. For instance, the concepts of Euler characteristic and Euler entropy have been introduced to a number of exactly solvable models [7, 10, 18–20] whose energy is described by a Hamiltonian function. Remarkably, in those systems the singularities of the Euler entropy were found to take place exactly at the transitions. This result resembles the statement of the Yang-Lee theorem [67], according to which the singular behavior exhibited by thermodynamic quantities in equilibrium phase transitions coincide precisely with the zeros of the system’s partition function.

The picture gains in complexity when it turns to the case of topological phase transitions occurring in complex networks. In this context, the Hamiltonian (energy) function is usually missing (or nonexistent) and instead of studying the system behavior as a function of energy, intrinsic correlations between the system constituents determined from empirical data define the network topology. Therefore, the Euler characteristic and entropy, as well as the Betti numbers (see text), emerge in this scenario as natural quantities to look into topological phase transitions in complex networks, particularly in the functional brain networks investigated here.

A number of computational topology tools has been successfully applied to the multidimensional analysis of complex networks [6]. For instance, persistent homology [14] has been employed across fields, such as contagion maps [68] and materials science [69]. In neuroscience, it has also yielded quite impactful results [31, 33, 37, 57, 70–72]. In this sense, the Blue Brain Project recently provided persuasive support based both on empirical data and theoretical insights for the hypothesis that the brain network comprises topological structures in up to eleven dimensions [51].

At this point one final remark is in order. Why is the above discussion about the interdisciplinary relations between topology, geometry, physics, neuroscience, and big data research important? The results of this work entail that, to a certain extent, correlations in functional brain networks are driven by principles that find some equivalence in the connections between phase transitions in physics, topology, and differential geometry. In this sense, one may argue that such relations might not be exclusive of brain networks, but instead can be part of a much wider picture featuring general data-driven complex systems that usually lack a Hamiltonian formulation. If so, the present study can shed light on a possible theoretical route based on such connections to explain the huge success of modelling similarity data using artificial intelligence and machine learning techniques.

### B. TDA approach and brain correlation matrices

Here we provide further technical details on the TDA approach applied to the functional brain networks and Erdős-Rényi random networks.

In the case of brain networks, we analyzed two datasets (HCP and VUmc, see below) containing information on the correlations among brain regions in the form of raw data and z-score data, respectively. These distinct forms were purposefully chosen to represent the diversity of approaches in the literature for measures of similarity among brain regions in brain networks [16].

We start with the analysis of the rs-fMRI measurements from the HCP dataset [73] under the 1000 Functional Connectomes Project [74]. It comprises data from 986 individuals, with the brain subdivided into *N* = 177 regions. The rs-fMRI measurements generate one time series for the activity of each particular brain region of each individual. It is thus possible to calculate the raw correlation between each pair of brain regions *i* and *j* of each individual by means of a Pearson correlation coefficient *C_ij_* ∈ [−1, 1], with *i, j* =1*,…, N*. Therefore, in this case one is left with an *N* × *N* matrix *C_ij_* of raw correlations among brain regions for each individual.

Once the correlation matrices *C_ij_* are built, the next step is the filtration process. In a physical system, where the energy is described by a Hamitonian function, the study of the topology of the configuration space can be done by sweeping up the energy levels in the equipotential energy surface [7, 10, 18–20] (see Fig. 1). Here, instead of the energy of a physical system or the variable that controls the height function in Morse theory, we define a filtration parameter in the form of a correlation threshold level *ε* as follows.

It is usually considered [75] that a small, nonzero value of *C_ij_* may just reflect noise, rather than the existence of any actual functional connection between brain regions *i* and *j*. To circumvent this problem, a thresholding is typically applied [16] in order to retain only correlations whose absolute value |*C_ij_*| lies above some given correlation threshold level *ε* ∈ [0, 1]. In mathematical terms, we write

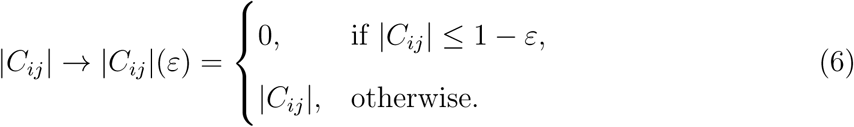

Conventionally, the diagonal elements |*C_ii_*| are set up to zero [75]. A functional brain network can thus be constructed from the matrix |*C_ij_*|(*ε*) in Eq. (6) by assigning an edge connecting the brain regions *i* and *j* if |*C_ij_*| ≠ 0 at the given value *ε*. In this sense, Eq. (6) defines a connectome matrix [16] of the functional brain network for each *ε*. For example, a network with no links corresponds to *ε* = 0, whereas a fully connected structure arises for *ε* = 1. Furthermore, since the edges are generally associated with values 0 *<* |*C_ij_*| ≤ 1, then the matrix also relates to the normalized weighted adjacency matrix in graph theory [16], whose elements represent the connection weights between each pair of nodes in the network.

Although thresholding is a widely used technique in functional brain networks, it is in fact not yet clear what is the best strategy to choose the proper threshold value [16]. At this point, an analogy with Morse theory can be useful since one could track the topological evolution of the brain network as a function of *ε* in a way similar to level sets in a process called filtration [14, 17, 71], as explained below.

A graph with nodes and linking edges can represent a network structure. The filtration of a complex network is obtained by thresholding its connectome matrix for values *ε* ∈ [0, 1] and subsequently ordering the resulting evolving graphs for increasing 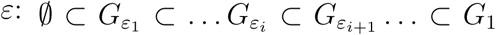, with the totally unconnected (empty) graph 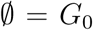, fully connected (complete) graph *G*_1_, and *ε_i_ < ε_i_*_+1_ (see Fig. 1). Here we implemented a filtration process using the Python NetwokrX library [76].

The bridge between TDA, network theory, and the topological approach to phase transitions is built through the calculation of topological quantities [17] such as the Euler characteristic and entropy and the set of Betti numbers as a function of *ε*. To this aim, it is necessary to compute the numbers of *k*-cliques in a graph, i.e., subgraphs with *k* all-to-all connected nodes (see Section 1.1). The computational effort to find a *k*-clique in a complex network is an NP-complete problem [77]. It means that, as more edges are attached, the numerical difficulty increases exponentially. For this reason, in the functional brain networks analyzed here we calculated the Euler characteristic using the NetworkX package up to the maximum threshold level *ε* =0.60. Moreover, we further observed that the convergence times to finish the computation of the *k*-cliques distribution are distinct for each individual brain network. Thus, as some networks presented prohibitively long convergence, we also applied a maximum cutoff processing time. Under this procedure, the brain networks of 420 individuals could reach the maximum threshold *ε* =0.60, see Figs. 3 and 4.

An even higher difficulty appeared in the calculation of the Betti numbers *β_n_* (see Section 1.1). In this case, in order to circumvent the long processing times we employed the concept of masked arrays [78], which performs averages of Betti numbers according to the number of brain networks that reached a given threshold level. Since the convergence time becomes increasingly longer for higher *ε*’s, not all networks reached the maximum value *ε* =0.50 displayed in Fig. 4, which explains the higher fluctuations in *β_n_* near this threshold level. We managed to compute, for a maximum period of 30 days of computing time, 712 individuals networks up to *ε* =0.50, which is above the first two topological phase transitions analyzed in detail here.

We now turn to the dataset from VU University Medical Center (VUmc dataset) [26], in which rs-fMRI measurements were performed using a different scanner and under different preprocessing steps in 15 control individuals with brain subdivided into *N* = 92 regions. Most numerical procedures applied to the VUmc dataset were quite similar to the ones of the HCP datset. Nevertheless, instead of raw correlation data, a different approach in terms of z-score values of correlations was employed in the VUmc dataset. Indeed, the matrix elements *C_ij_* were obtained as in the HCP dataset, in the form of a Pearson correlation coefficient. However, after taking their absolute value we computed the z-score values of correlations for each matrix element, i.e., the number of standard deviations by which the correlation value differs from the average, and proceeded the subsequent TDA study as in the case of the HCP dataset.

Lastly, we also mention that a similar analysis was performed in the Erdős-Rényi random networks, but with the probability *p* of linking two nodes playing the role of the correlation threshold level *ε* in the filtration process. In this case, however, we did not face the difficulty related to long processing times to compute the numbers of *k*-cliques since an exact analytical expression [35] is available for Erdős-Rényi graphs, see Eq. (1). Thus, we were able to sweep the whole interval *p* ∈ [0, 1] in the calculation of the Euler characteristic and entropy of Erdős-Rényi networks, as shown in Fig. 2. Nevertheless, for the computation of the Betti numbers in Fig. 2, since an analytic expression for *β_n_* is not at hand, we actually performed numerical simulation of smaller Erdős-Rényi networks (*N* = 25 nodes) and *p* ∈ [0, 0.7].

## Supporting information

Video - Topological Phase Transition

Interactive brain network figures

## V. ACKNOWLEDGEMENTS

F.A.N.S., E.P.R., M.D.C.F. and M.C. acknowledge financial support from CAPES, CNPq, and FACEPE through the PRONEX Program (Brazilian agencies). L.D. is funded by an NWO Veni (016.146.086) and a Branco Weiss Fellowship. Data were provided by the Human Connectome Project, WU-Minn Consortium (Principal Investigators: David Van Essen and Kamil Ugurbil; 1U54MH091657), funded by the 16 NIH Institutes and Centers that support the NIH Blueprint for Neuroscience Research, and by the McDonnell Center for Systems Neuroscience at Washington University. Data processed at VUmc were kindly provided by Dr. M. Raemaekers, from Dept. of Neurology and Neurosurgery, Brain Center Rudolf Magnus, University Medical Center Utrecht. F.A.N.S. would also like to thank fruitful discussions with Sostenes Lins, Heather Harrington, Ulrike Tillmann, Fernando Moraes, Eamonn A. Gaffney, Jeffrey Giansiracusa, Pawel Dlotko, Marco Baroni, Cleide Martins, Renê Rodrigues Montenegro Filho, Bertrand Berche, Victor Caldas, Dante Chialvo, Stephan Boettcher, and Tom Shimizu, during the development of this work.

## VI. AUTHOR CONTRIBUTIONS

F.A.N.S. developed the theoretical formalism and performed the calculations and numerical simulations of the Erdős-Rényi random networks and functional brain networks. L.D. and F.A.N.S. proposed the search for topological phase transitions in brain networks. E.P.R. and M.D.C.F. proceeded the scaling analysis and fitting of the data near the phase transition point. F.A.N.S. and E.P.R. wrote the first version of the manuscript. All authors were involved in the subsequent writing and development of the manuscript. All authors planned the research and discussed the results.

